# Inorganic sulfur fixation via a new homocysteine synthase allows yeast cells to cooperatively compensate for methionine auxotrophy

**DOI:** 10.1101/2022.03.14.484209

**Authors:** Jason S.L. Yu, Benjamin M. Heineike, Johannes Hartl, Clara Correia-Melo, Simran Kaur Aulakh, Andrea Lehmann, Oliver Lemke, Federica Agostini, Cory T. Lee, Vadim Demichev, Christoph B. Messner, Michael Mülleder, Markus Ralser

## Abstract

The assimilation, incorporation, and metabolism of sulfur is a fundamental process across all domains of life, yet how cells deal with varying sulfur availability is not well understood. We studied an unresolved conundrum of sulfur fixation in yeast, in which an organosulfur-auxotrophy caused by deletion of homocysteine synthase Met17p is overcome when cells are inoculated at high cell density. We discovered that an uncharacterized gene YLL058Wp, herein named Hydrogen sulfide utilizing-1 (*HSU1*), acts as a homocysteine synthase and allows the cells to substitute for Met17p by re-assimilating hydrosulfide ions leaked from *met17Δ* cells into O-acetyl-homoserine and forming homocysteine. Our results show that cells can cooperate to achieve sulfur fixation, indicating that the collective properties of microbial communities facilitate their basic metabolic capacity.

**Summary:** Sulfur limitation activates a dormant hydrogen sulfide fixation route via a novel homocysteine synthase Hsu1p (YLL058Wp).

## Introduction

Despite decades of effort, a large proportion of the coding sequences expressed from eukaryotic genomes are not or are incompletely functionally annotated (Haynes et al., 2018; Stoeger et al., 2018; Wood et al., 2019). One contributing factor to this situation is that many laboratory experiments are conducted under a limited set of standardized growth conditions that do not fully represent the range of conditions organisms are exposed to in nature (Hillenmeyer et al., 2008). The budding yeast *Saccharomyces cerevisiae* is a key model organism for the discovery and characterization of gene function in eukaryotes. Moreover, because of its importance in biotechnology, the response to varying carbon and nitrogen sources has been extensively studied in this organism (Lange & Heijnen, 2001; Zaman et al., 2008). However, although the utilization of sulfur is an equally fundamental process, far less is known beyond the core genes that are involved in the assimilation and utilization of inorganic sulfur into metabolically useful organosulfur compounds (Ghosh & Dam, 2009; Hébert et al., 2011; Muyzer & Stams, 2008).

In many eukaryotes, sulfur is primarily assimilated in the form of sulfate (SO_4_^2-^), which becomes successively reduced to sulfide (S^2-^). A key enzyme in the eukaryotic sulfur assimilation process is homocysteine synthase which in *S. cerevisiae*, is encoded by the *MET17* gene (also known as *MET15 or MET25*). Met17p catalyzes the fixation of inorganic sulfide with O-acetylhomoserine (OAHS) to form homocysteine (Fig. 1A). Homocysteine is subsequently converted to methionine via the methionine synthase (Met6p) or into cysteine via cystathionine-β-synthase and cystathionine-γ-lyase (Cys4p and Cys3p, respectively) (D’Andrea et al., 1987; Kerjan et al., 1986; Yamagata, 1981). Homocysteine therefore provides the central organosulfur pool from which the key sulfur-bearing amino acids methionine and cysteine are synthesized (Cherest & Surdin-Kerjan, 1992; Thomas & Surdin-Kerjan, 1997).

**Figure 1:**
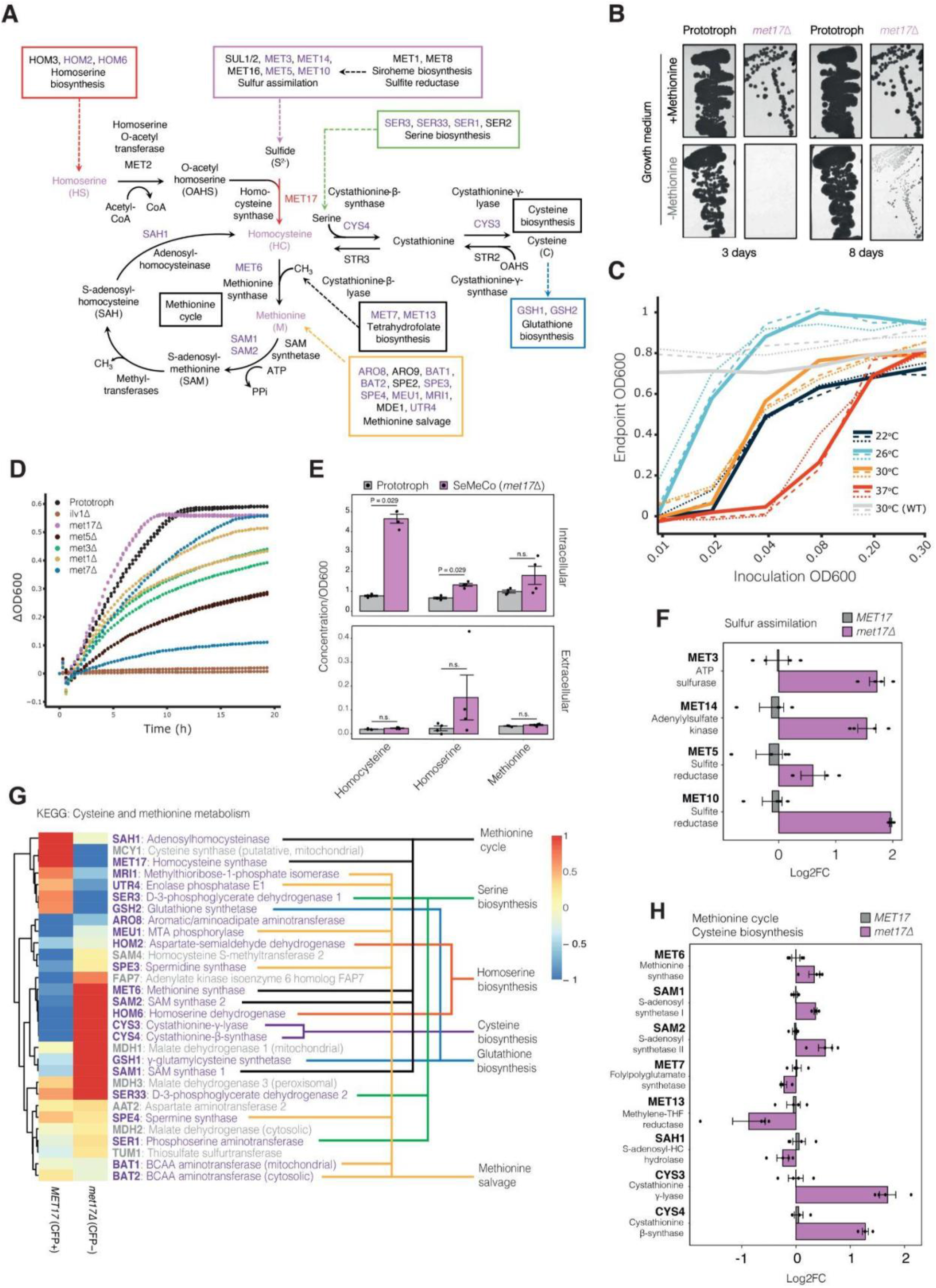
Characterisation of the *met17*Δ growth paradox by metabolomics and proteomics. **(A)** The core and associated metabolic pathways that feed into organosulfur biosynthesis centered around homocysteine, including the gene names that encode for the respective enzymes in *S. cerevisiae.* **(B)** Streak cultures of prototrophic wild-type and *met17*Δ strains on agar with and without methionine, imaged at 3- and 8-days incubation at 30°C. **(C)** Liquid cultures of *met17*Δ strain in minimal media without methionine inoculated at 6 different cell densities (OD_600_ = 0.01 to 0.3) and cultured at 4 different temperatures (22, 26, 30, 37°C) for 48h. Gray indicates prototrophic wild-type control strain cultured at 30°C. Lines indicate three replicates per condition. **(D)** Growth of methionine auxotrophic deletion strains with prototrophic and *ilv1*Δ as control strains in minimal medium. All strains have a background of *met17*Δ, utilizing intracellular stores of organosulfur compounds to maintain their growth. **(E)** Metabolomic analysis and measurement of homocysteine, homoserine and methionine in extracellular and intracellular fractions between exponentially growing prototrophic and SeMeCo cultures normalized to the final OD. Data is from 2 biologically independent replicates containing 4 technical replicates (n=8). Adjusted p-values from unpaired Wilcoxon rank sum test are as indicated, n.s. indicates no statistical significance. **(F)** Differential protein expression analysis of sulfur assimilation enzymes between sorted CFP+ (*MET17*) and CFP- (*met17*㥄), where n=4 biologically independent replicates. **(G)** KEGG pathway analysis and hierarchical clustering of sorted cells as in (F) of all detected enzymes involved in cysteine and methionine metabolism characterized by associated metabolic pathways (colored). (**H)** Differential protein expression analysis of core enzymes between sorted CFP+ (*MET17*) and CFP- (*met17*Δ) involved in the methionine cycle and cysteine biosynthesis where n=4 biologically independent replicates.

Deletion of *MET17* (*met17*Δ) in budding yeast renders cells auxotrophic for organosulfur compounds such as methionine, cysteine, homocysteine or S-adenosyl methionine (Masselot & de Robichon-Szulmajster, 1975; Singh & Sherman, 1974; Thomas & Surdin-Kerjan, 1997; Yamagata, 1976). This led to the widespread use of the *met17*Δ allele as an auxotrophic marker for genetic experiments (Cost & Boeke, 1996). *met17*Δ alleles were crossed into many laboratory strains, including the W303 and S288c derivatives that gave rise to the yeast deletion collection (Winzeler et al., 1999). However, this led to reports of an atypical growth phenotype. When thick patches of *met17*Δ cells are replicated, they can overcome the auxotrophy and continue growth in the absence of an organosulfur source. The phenomenon reportedly occurs in a temperature-sensitive manner, with growth of the auxotrophs being most evident at 22°C and decreasing linearly with temperature up to 37°C (G. J. Cost, 1999). This puzzling cell-density and temperature-sensitive phenotype has also been described in distantly related yeasts (i.e. *Candida albicans* and *Yarrowia lipolytica*) (Edwards et al., 2020; Viaene et al., 2000) suggesting it is not a species-specific phenomenon. Among the possible explanations for this paradoxical phenotype, Cost and Boeke (G. J. Cost, 1999) suggested that *met17*Δ could leak and share organosulfur metabolites between cells once they reached a critical cell density.

Here, we studied the biochemical basis of the *met17*Δ phenotype and discovered a previously overlooked metabolic bypass that explains the growth of *met17*Δ in absence of organosulfur compounds *(met17*Δ paradox). We found that an uncharacterized protein, which we herein have named Hydrogen Sulfide Utilizing-1 (Hsu1p, formerly YLL058W), encodes a metabolic enzyme that directly assimilates sulfur from hydrogen sulfide, thereby enabling cell growth by resolving methionine auxotrophy once critical concentrations of sulfide are leaked from *met17*Δ cells. We demonstrate that Hsu1p functions as a homocysteine synthase, performing the fixation of inorganic sulfide in place of Met17p, thereby generating the homocysteine required for methionine and cysteine biosynthesis.

## Results

### The conditional nature of organosulfur auxotrophy in *met17*Δ yeast

The ability of *met17*Δ yeast strains to overcome organosulfur auxotrophy at high cell density was described by Cost and Boeke (Cost & Boeke, 1996; G. J. Cost, 1999). This phenotype is non-heritable, which ruled out adaptive, secondary mutations concurrent with *met17*Δ as the cause. We studied the phenotype in two yeast strains in the haploid lab strain BY4741 background (Brachmann et al., 1998): i) a prototrophic strain in which auxotrophies were repaired by genomic integration of the missing genes (*HIS3, LEU2, URA3, MET17)*, and ii) an analogous strain in which the *met17*Δ allele was not repaired (*HIS3, LEU2, URA3, met17*Δ). Prototrophy was restored in this background to enable the use of minimal (non-amino acid supplemented) media for the study of amino acid biosynthesis (Mülleder et al., 2012). Confirming the robustness of the unusual growth phenotype (Cost & Boeke, 1996; G. J. Cost, 1999), upon low density inoculation, the *met17*Δ strain did not grow when methionine was absent from the in minimal medium. However, when replicated as dense patches, the *met17*Δ strain was able to grow on this medium over the course of 8 days (Fig. 1B).

We started our investigations into the biochemical basis of this phenotype by ruling out amino acid contaminations in commercial yeast nitrogen broth (YNB), as a common source of conflicting and unusual growth phenotypes. We validated the concentration of sulfur containing amino acids using liquid chromatography-selected reaction monitoring (LC-SRM) (Mülleder et al., 2017). We detected levels of methionine, homocysteine and glutathione at a concentration at least 100,000-fold lower than typical levels found in replete yeast media (Fig. S1A). Therefore, the growth of *met17*Δ cells in the absence of methionine was not due to the presence of growth-relevant concentrations of sulfur containing amino acids in the growth medium. In order to have a more experimentally tractable system to further our investigations, we then explored the conditions, under which the growth phenotype was reproducible in liquid culture. We inoculated the *met17*Δ strain into methionine-free medium at a range of initial densities (OD_600_ = 0.01 to 0.3) and cultured under four temperatures (22, 26, 30 and 37°C), recording the endpoint OD_600_ at 24h (Fig. S1B) and 48h (Fig. 1C) as a measure of growth. At all temperatures tested growth could only be observed with high-density inoculations (OD_600_ = 0.02 to 0.3), in agreement with previous observations (Cost & Boeke, 1996). In contrast, wild-type prototrophic strains could robustly grow at all inoculation densities at 30°C (Fig. 1C). Although we did not observe a strict correlation of temperature with growth as previously reported, growth of low-density inoculations (OD_600_ = 0.02 to 0.03) in the absence of methionine was greatly enhanced at a lower temperature than typically used during the cultivation of lab strains (26°C compared with 30°C by 48h). However, growth of the *met17*Δ strain was never observed under any condition when the initial inoculation density was lower than 0.01. The same density and temperature conditions permissive for growth were also confirmed in larger scale cultures (Fig. S1C), confirming that the ability to overcome organosulfur auxotrophy is truly dependent on cell density opposed to culture size.

Next, to probe whether the cell density limitation is specific to *met17*Δ or whether it is a more general phenomenon related to methionine or amino acid auxotrophy, we compared the growth of the *met17*Δ strain to other enzyme knock-out strains of sulfur assimilation pathways (*met1*Δ, *met3*Δ, *met5*Δ or *met7*Δ) or amino acid metabolism (*argΔ, hisΔ, trpΔ, aroΔ, lysΔ, phe*Δ), including *ilv1*Δ, as a *bona fide* auxotrophic control (Kakar & Wagner, 1964; Thuriaux et al., 1971). All strains except *met17*Δ were growth deficient, despite high-density inoculation and the absence of amino acid supplementation (Fig. 1D, Fig. S1D) This result suggested that the ability to overcome the growth defect is not a common phenotype of methionine auxotrophs but rather a specific phenotype that occurs upon the deletion of *MET17.* We speculated that either another enzyme replaces the function of Met17p in these conditions, or else a metabolic shunt exists that allows yeast to circumvent the canonical sulfur utilization pathway to fuel growth.

### The ability to overcome organosulfur auxotrophy does not require the exchange of homocysteine or methionine

Cost and Boeke (G. J. Cost, 1999) speculated that *met17*Δ cells could overcome the growth defect through the sharing of organosulfur metabolites between cells to growth-relevant levels. Indeed, previous work from us and others has shown that communal yeast cells can effectively share metabolites to overcome auxotrophies (Campbell et al., 2015; Ponomarova et al., 2017). We explored self-establishing metabolically cooperating (SeMeCo) communities, a synthetic yeast community designed to study metabolite exchange interactions between auxotrophs (Campbell et al., 2015). SeMeCo communities are composed of cells that express different combinations of auxotrophic markers, and establish from a prototrophic founder cell through the stochastic loss of plasmids encoding one or more auxotrophic markers that compensate for auxotrophies present in the genome. In the BY4741 background, these are *met17Δ, his3Δ, leu2Δ* and *ura3Δ*, which induce methionine, histidine, leucine and uracil auxotrophy respectively. In SeMeCos, the degree of metabolite sharing influences the frequency of a respective auxotroph in the population. Examining data recently acquired by us revealed that *met17*Δ cells are the most dominant subpopulation within exponentially growing SeMeCo cultures (~60%), and more frequent than the three other auxotrophies (Fig. S1E). (Yu et al., 2022) This suggested that metabolite sharing is more effective at overcoming the loss of *MET17*, at least when compared to the other three other auxotrophies. In order to understand the role that metabolite sharing plays in overcoming the loss of *MET17* in the SeMeCo system, we recorded the metabolic profiles of SeMeCos by LC-SRM and compared these against the metabolic profiles of prototrophic communities. In SeMeCo communities dominated by *met17*Δ cells, we detected a relative increase in intracellular homocysteine and homoserine concentrations compared to prototrophic communities (Fig. 1E, intracellular). However, extracellular concentrations of homocysteine and methionine were not statistically different, although homoserine levels were elevated upon the exclusion of an outlying measurement (Fig. 1E, extracellular). This result suggested that indeed, the *met17*Δ auxotrophy might be overcome through the exchange of an upstream precursor rather than directly via methionine sharing.

### *met17*Δ cells upregulate enzymes involved in sulfur assimilation, cysteine and methionine metabolism

We have previously generated proteomes from sorted auxotrophs and prototrophs from SeMeCo cultures (Yu et al., 2022). To distinguish auxotrophs from prototrophs, we used cosegregation of cyan fluorescent protein (CFP) with the relevant auxotrophic marker expressed from the same plasmid (PRIDE project: PXD031160). A differential expression analysis comparing specifically the *MET17* and *met17*Δ subpopulations revealed a clear upregulation of sulfur assimilation enzymes (Met3p, Met14p, Met5p and Met10p) and enzymes of the methionine salvage pathway (Meu1p, Spe3p, Spe4p, Bat1p and Bat2p) in the *met17*Δ background. This result was indicative of a feedback response to the lack of methionine via increased levels of sulfate assimilation or sulfur recycling (Fig. 1F, Fig. 1G). Counterintuitively, several of the downstream enzymes, including Met6p and Gsh1p, that should carry no flux in the absence of Met17p, were increased in their expression level (Fig. 1A, Fig. 1G, Fig. 1H). In addition, the abundance of other core enzymes of the methionine cycle (Met6p, Sam1p, Sam2p), serine (Ser1p, Ser33p), and cysteine (Cys3p, Cys4p) biosynthetic pathways, were also increased (Fig. 1H). Together, we hence observed that when the growth defect of a *met17*Δ is overcome in SeMeCo communities, one observes an increase in intracellular homocysteine and an increased expression of downstream enzymes. This data argued that homocysteine is further metabolized despite the absence of Met17p, suggesting the existence of a metabolic bypass to the homocysteine synthase activity of Met17p (Fig. 1G).

### Hydrogen sulfide fixation drives growth in the absence of methionine

We speculated that the intracellular accumulation of homocysteine in *met17*Δ cells (Fig. 1E) could indicate the presence of an enzyme-catalyzed reaction that forms homocysteine independent of Met17p. As sulfide overflow is a defining characteristic of *met17*Δ cells (Cost & Boeke, 1996; G. J. Cost, 1999), one possible explanation would be the presence of an enzyme that can utilize sulfide or its protonated forms (hydrosulfide (HS^-^) and hydrogen sulfide (H_2_S)) to incorporate sulfur and form homocysteine (Cost & Boeke, 1996; G. J. Cost, 1999). To test this hypothesis, we exploited the cell density dependency of *met17*Δ cells in liquid culture that leads to the resolution of organosulfur auxotrophy as a readout (Fig. 1C, Fig. S1B,C). When conditioned medium from *met17*Δ cells cultured either at high-density (OD_600_ = 0.08) or low-density (OD_600_ = 0.01) was filtered and inoculated with fresh cells at low-density, only the filtered medium from the high-density culture could support cell growth. Moreover, the ability to support growth was abolished upon a lead acetate precipitation of H_2_S (Fig. 2A) (*14, 16*). To further substantiate whether H_2_S could be responsible for the growth, we supplemented two low-density cultures with 0.2 mM sodium hydrosulfide (NaHS) that dissociates to form hydrosulfide ions, and subsequently, both aqueous and gaseous forms of H_2_S (Fig. 2B, reactions (ii-iv)). Taking advantage of the phase equilibria of H_2_S, one culture was left unsealed to drive the equilibria towards the formation and dissipation of gaseous H_2_S from the system (Fig. 2B, reaction (iv)). Growth was only observed in the sealed culture, consistent with the idea that the concentration of H_2_S is required to overcome the *met17*Δ phenotype (Fig. 2C). In parallel, we manipulated the phase equilibria in the opposite direction, allowing the accumulation of aqueous H_2_S in low-density cultures from outgassing from either a high-density culture or a 20 mM solution of NaHS via connecting the vessels with a small piece of rubber tubing. In both situations, growth was promoted in the low-density *met17*Δ culture (Fig. 2D). To further substantiate that it is H2S which allows the overcoming of the auxotrophy, we evaluated the presence and uptake of H_2_S in the different cultures and growth conditions exploiting 7-azido-4-methylcoumarin (AzMC). In the presence of H_2_S, AzMC undergoes selective reduction of the azido moiety to form 7-amino-4-methylcoumarin (AMC) which fluoresces when exposed to ultraviolet light (Thorson et al., 2013). When minimal media was compared with cultures inoculated at low-density, no significant difference in AMC fluorescence was observed. Conversely, the fluorescence in NaHS supplemented cultures decreased in the presence of cells, suggesting utilization of H_2_S and its concomitant depletion from the medium (Fig. 2E). This fluorescence assay indicated that high-density cultures had higher levels of aqueous H_2_S, relative to low-density cultures (Fig. 2A). The presence of higher levels of aqueous H_2_S in high-density cultures was orthogonally confirmed by the precipitation of lead sulfide in high-density cultures than in low-density cultures (Fig. 2E).

**Figure 2:**
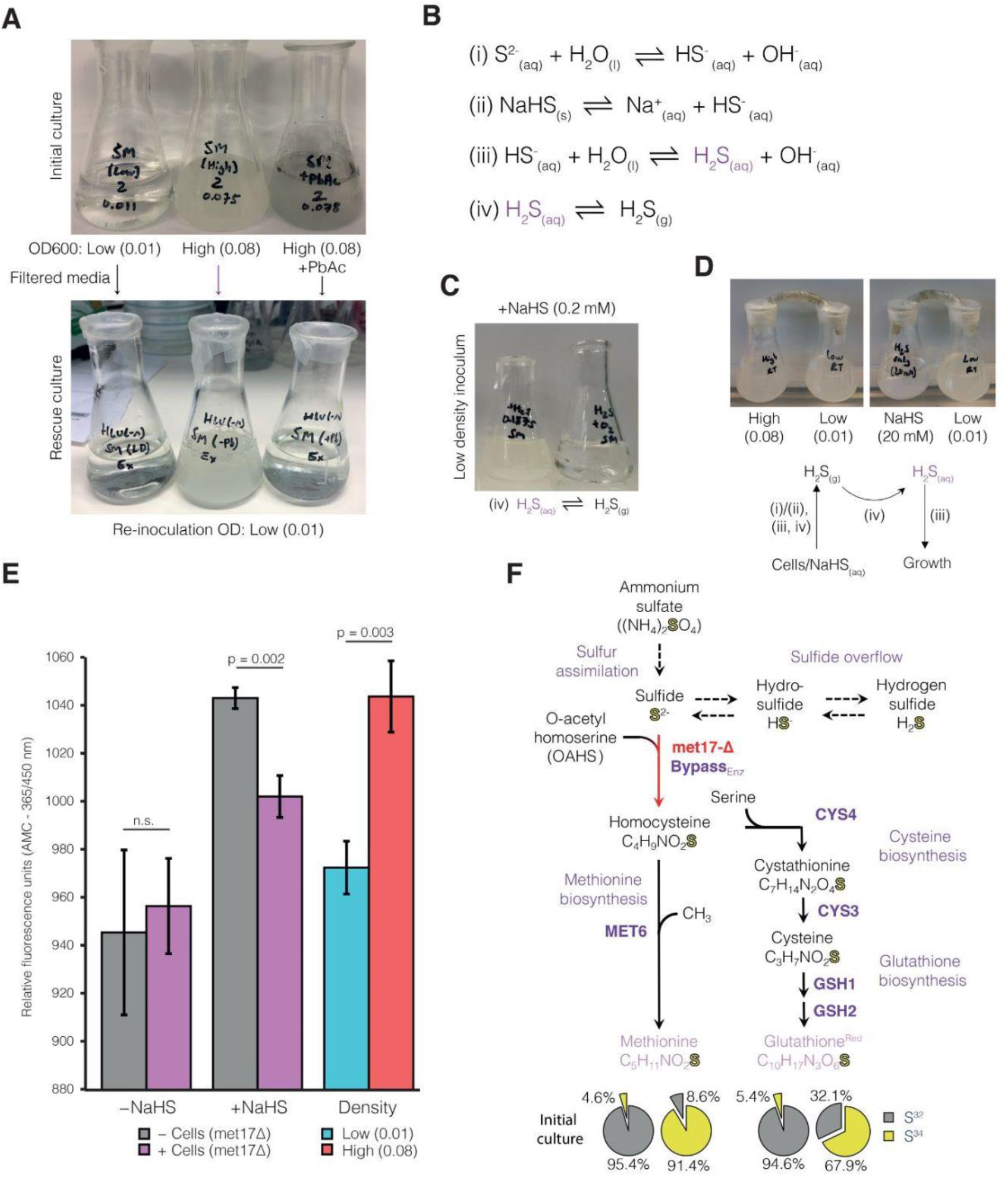
YLL058Wp utilizes hydrogen sulfide to rescue organosulfur auxotrophy in *met17*Δ strains. **(A)** *met17*Δ cultures inoculated at low starting density (OD_600_ = 0.01) and high densities (OD_600_ = 0.08) (upper panel, left and middle). One flask of high density inoculation was supplemented with lead (II) acetate to precipitate H_2_S (upper panel, right). Conditioned media from these three cultures was subsequently filtered, replenished with yeast nitrogen broth (YNB) and glucose and re-inoculated at low-density (OD_600_ = 0.01) (lower panel). Data are representative from two biologically independent experiments. **(B)** Reaction schemes. Hydrolysis of sulfide is in equilibrium with hydrosulfide and hydrogen sulfide (i, iii, iv). NaHS can serve as a hydrosulfide donor (ii) in solution. Hydrogen sulfide preferentially undergoes phase transition to a gaseous state (iv) in unsealed vessels or establishes a dynamic equilibrium with its aqueous counterpart in sealed vessels. **(C)** Growth of *met17*Δ cultures supplemented with NaHS and inoculated at low-density, sealed vs unsealed. **(D)** Cultures inoculated with the *met17*Δ strain where the vessels are connected via rubber tubing to facilitate gaseous exchange, either between a high-density culture or a high concentration solution of NaHS (20 mM). Schematic indicates the sulfur chemistry occurring at each stage to facilitate the growth of low-density cultures. **(E)** Quantification of hydrogen sulfide production and uptake via conversion of AzMC to AMC (*21*) between unsupplemented and NaHS supplemented cultures inoculated at low density in the presence (purple) or absence (gray) of *met17*Δ cells. Blue and red bars indicate a separate experiment where hydrogen sulfide production was quantified *met17*Δ cultures inoculated at either low (blue) or high (red) density, in a similar experiment as shown in Fig. 2A. p-values calculated via two-tailed Student’s *t*-test, where n=3 independent replicates. **(F)** Isotopic tracing experiment with pathway map indicating the biosynthetic transfer of isotopic S^34^ from ammonium sulfate to methionine (left group) and glutathione (right group) via methionine and cysteine biosynthetic pathways respectively. Pie charts indicate the percentage of S^34^ for cultures supplemented with non-labelled ammonium sulfate (left chart per group) or when supplemented with S^34^-labeled ammonium sulfate (right chart per group).

Finally, to test if sulfur does indeed transition from sulfate through to the biosynthesis of cysteine and methionine in the absence of *MET17*, we performed isotopic tracing with S^34^-labeled ammonium sulfate (Fig. 2F, Fig. S2A). Uptake of this isotope label by sulfur assimilation via the upregulation of the associated enzymes should lead to S^34^ dissemination into all subsequent sulfur bearing derivatives via sulfide overflow (Fig. 1A, Fig. S2B). We quantified the ratios of S^32^ and S^34^ sulfur in methionine and reduced glutathione, the downstream products derived from homocysteine and cysteine biosynthesis, respectively, using liquid chromatography mass spectrometry (LC-MS). When native (S^32^) ammonium sulfate was supplied, the assay detected S^34^ at its expected environmental iso topological abundance (4-6%). Conversely, when we supplied S^34^-labeled ammonium sulfate, S^34^-methionine (91.4%) and S^34^-reduced glutathione (67.9%) accumulated to levels above natural abundances, indicative of sulfur transfer from inorganic sulfate to organic sulfur-bearing metabolites. Thus, *met17*Δ assimilated inorganic sulfur from ammonium sulfate, and yeast cells are competent in the use of H_2_S as a source for methionine and cysteine biosynthesis.

### A targeted genetic screen identifies the bypass enzyme responsible for H_2_S-mediated growth in the absence of methionine

Having established homocysteine formation via H_2_S fixation as the process by which methionine and cysteine auxotrophy could be overcome, we next used a targeted genetic approach to identify the enzyme responsible. We started by identifying strains in the yeast deletion collection (Winzeler et al., 1999) that are associated with sulfur metabolism. We selected 15 strains (Table S1) that carried a deletion in the sulfur metabolism-associated gene in addition to deletions of four auxotrophic markers in their genome that include *MET17* (*his3*Δ*leu2*Δ*ura3*Δ*met17*Δ) based on two criteria: i) the deleted gene was a part of either the sulfur assimilation or organosulfur biosynthetic pathways, or ii) the deleted gene had homology to Met17p according to the eggNOG database (Baudin et al., 1993; Shoemaker et al., 1996) (Fig. 1A). These selected deletion strains were cultured in synthetic drop-in media and tested for *met17*Δ growth in the presence of 0.4 mM NaHS. Mutant strains lacking enzymes directly upstream of Met17p (Sul1/2p, Met3p, Met14p, Met5p and Met10p) demonstrated robust growth only in the presence of NaHS, whilst the strain lacking the methionine synthase Met6p was unable to grow. These growth phenotypes indicated that H_2_S utilization and the resolution of organosulfur auxotrophy in the presence of NaHS was independent of sulfur assimilation, but dependent on methionine biosynthesis (Fig. 3A, Fig. S2B). Critically, these experiments also revealed the dependency of the H_2_S utilization pathway on Met2p, the homoserine-O-acetyltransferase that is responsible for the conversion of homoserine to O-acetylhomoserine (OAHS), a substrate shared by both Met17p and Str2p, implying a similar catalytic mechanism (Fig. 3B, Fig 1A, (Masselot & de Robichon-Szulmajster, 1975)). Of the strains deficient in enzymes homologous to Met17p, the *met17Δcys3Δ* strain also could be rescued by NaHS supplementation, which suggests the existence of a direct route towards cysteine formation via H_2_S, possibly via cysteine synthase activity (Fig. S2C, Fig. 3B, gray pathway,(Cherest & Surdin-Kerjan, 1992; B. Ono et al., 1991, 1992)). Notably, loss of cystathionine-γ-synthase and cystathionine-β-lyase (Str2p and Str3p), two other Met17p homologs, also led to a complete and partial loss of growth rescue by NaHS supplementation respectively. This suggested that the degree of transsulfuration from cysteine to homocysteine may act to regulate the efficiency of the bypass reaction, or that Str2p might itself be the bypass enzyme. Intriguingly, we observed that deletion of an uncharacterized Met17p homolog, YLL058Wp, also led to the loss of the ability to utilize H_2_S for growth (Fig. 3A). This defect was not observed upon deletion of the remaining Met17p homologs Irc7p, YHR112Cp, and YML082Wp (Fig. S2C), indicating that YLL058Wp could also potentially function as the bypass enzyme. We next asked if the gene products of *STR2* and *YLL058W* could therefore operate as a homocysteine synthase in place of Met17p to rescue organosulfur auxotrophy. Str2p has been shown to catalyze the conversion of cysteine to cystathionine using OAHS as a substrate and, operating in tandem with Str3p, permits the transsulfuration of cysteine to methionine (Cherest & Surdin-Kerjan, 1992; Hansen & Johannesen, 2000). Since Str2p shares the same substrate as Met17p, there was a possibility it could incorporate sulfur via a secondary or promiscuous reaction. The catalytic properties of YLL058Wp, our other candidate, have not previously been experimentally characterized.

**Figure 3:**
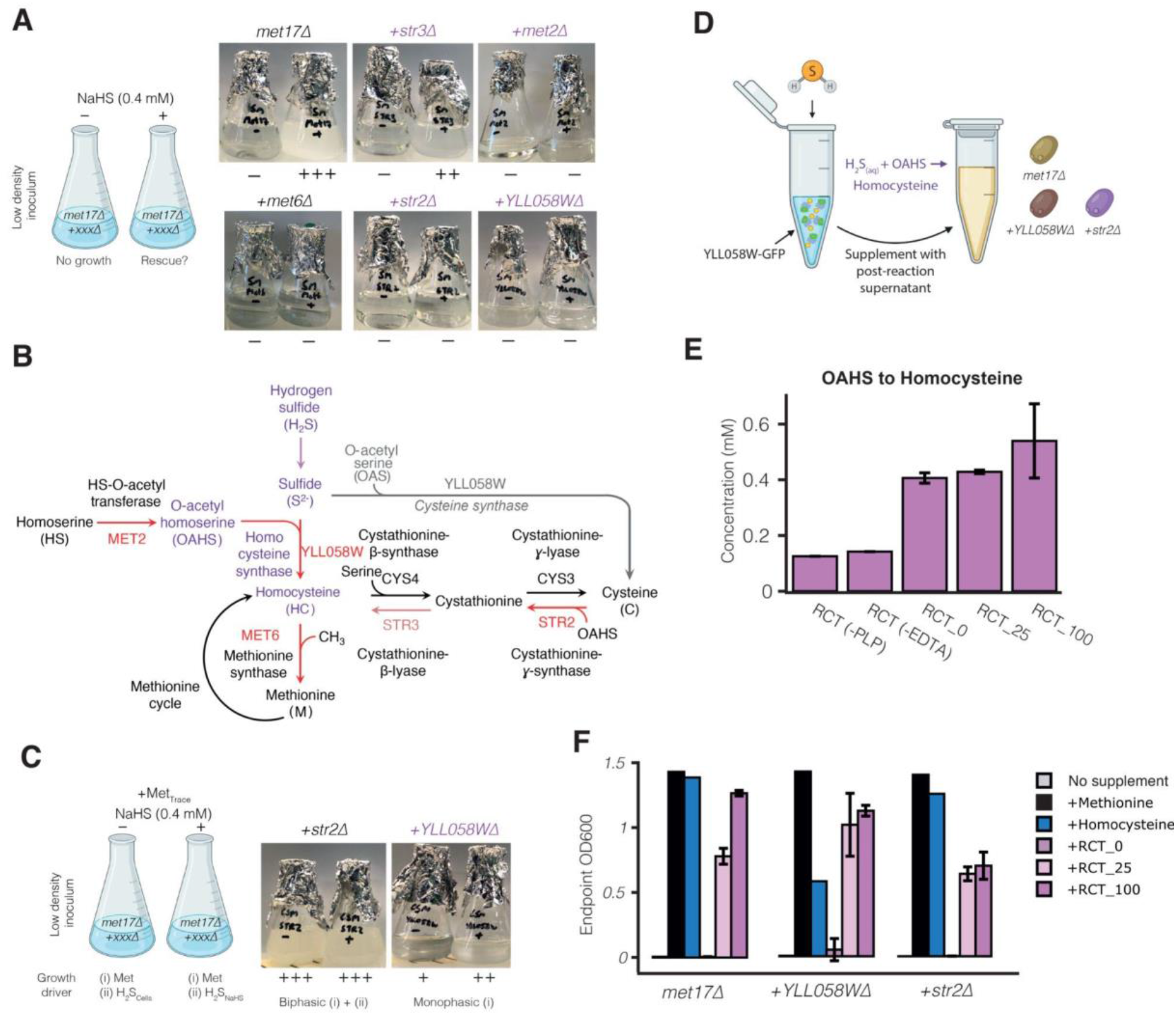
Identification and characterisation of YLL058Wp as the homocysteine synthase responsible for hydrogen sulfide fixation. **(A)** A targeted genetic screen of double-knockout strains lacking, in addition to *met17*Δ, the genes encoding the homoserine (Met2p), methionine (Met6p) biosynthesis pathways and selected orthologs of Met17p (Str3p, Str2p, YLL058Wp). Cultures were inoculated at low-density either in the presence or absence of NaHS which rescues growth in *met17*Δ strains. A mutant strain carrying *met17*Δ alone was used as control. Complete or partial loss of NaHS rescue phenotype in these cultures indicate involvement of the deleted enzyme in the metabolic bypass (purple). Images are representative of 3 independent biological replicates. **(B)** Summary of the deletion mutants that result in a loss (red) or partial loss (pink) of the NaHS-mediated growth rescue mapped onto a theoretical bypass pathway. Purple indicates key metabolites and enzyme activities involved in the bypass. Gray pathway indicates the conversion from OAS to cysteine that we observe *in vitro* with YLL058Wp, but which is unlikely to exist in vivo in an S288C background (see text). **(C)** Methionine limitation screen comparing *met17*Δ*str2*Δ and *met17*Δ*YLL058W*Δ. Low amounts of methionine were provided to fuel the first phase of growth (monophasic), allowing cells to reach a critical density at which sufficient hydrogen sulfide leaked from cells to allow further growth (biphasic). In this scenario, NaHS supplementation enables the second phase of growth to proceed if the bypass is functional by allowing cells to freely use H_2_S as a sulfur source regardless of limited concentrations of methionine. **(D)** Schematic indicating the setup of *in vitro* enzymatic reaction and growth assays used to assess the ability of OAHS reaction supernatants to rescue the indicated deletion strains (*met17*Δ, *met17*Δ*YLL058W*Δ, *met17*Δ*str2*Δ). **(E)** Quantification of thiol concentrations via Ellman’s reagent and spectrometry using post-reaction supernatants where immunoprecipitated YLL058W-GFP was incubated with OAHS as the organic substrate. RCT_0, 25 and 100 indicate volumes of enzyme-resin slurry used in the reaction. RCT(-PLP or -EDTA) indicates reactions whereby pyridoxal-5’ phosphate (PLP) or EDTA has been omitted from the reaction buffer. Yield = [product]/[substrate], OAHS: 0.54/5 = 11%. Data are mean thiol concentrations ± SD where n = 3 biologically independent replicates. **(F)** Growth assays whereby each strain was inoculated at low-density and supplemented with 50 μl of the indicated *in vitro* reaction product (n=3). Final OD_600_ measured following 5 days of outgrowth. For OAHS reaction supplements, data are mean OD_600_ ± SD where n = 3 biologically independent replicates. OD_600_ reached in minimal media and supplementation with homocysteine or methionine (single cultures each) are shown for comparison. Schemes for Fig. A, C and D were created with BioRender.com.

In order to characterize how these two enzymes contributed to the bypass reaction, we devised an experiment in which we supplemented only a limited concentration of methionine into minimal media that was rapidly depleted (Fig. 3C). In essence, the assay separates growth into two phases, the first phase is driven by methionine until the cells have reached the critical cell density that is required by the cells to overcome the *MET17* deficiency using the unknown bypass enzyme. The *met17*Δ*str2*Δ strain maintained robust growth, both in the presence and absence of supplemented NaHS, indicating that the bypass reaction does not require Str2p. In contrast, growth of the *met17*Δ*YLL058W*Δ strain ceased after the initial biomass accumulation phase that is driven by the presence of methionine, and did not achieve robust growth even in the presence of ample NaHS (Fig. 3C). Thus, YLL058Wp is necessary and sufficient for the cell to assimilate sulfide, whilst Str2p appears to indirectly affect the bypass reaction.

### YLL058Wp is a homocysteine and cysteine synthase that fixes sulfur from hydrogen sulfide

Having identified an enzyme required for the bypass reaction, we next sought to further characterize the expression and activity of YLL058Wp, using a yeast strain that expresses a YLL058W-GFP fusion protein in an auxotrophic background lacking Met17p (*leu2*Δ*ura3*Δ*met17*Δ). We first independently verified that the open reading frame was intact by PCR (Fig. S3A (Huh et al., 2003)). Expression of the GFP fusion protein was poor when cultured in rich media such as YPD. The expression of YLL058W-GFP increased 1.5-fold in minimal media containing leucine, uracil and NaHS (SM+LU+NaHS), (Fig. S3B). We next immunoprecipitated YLL058Wp-GFP from yeast cells grown under these conditions and tested *in vitro* if the enriched protein was capable of transferring the sulfur from H_2_S onto OAHS to form homocysteine (Fig. 3D, Fig. S3C). We also tested the ability of the enzyme to function as a cysteine synthase using O-acetylserine (OAS) as an alternative substrate, since this could explain the growth of the *met17*Δ*cys3*Δ strain in the presence of NaHS (Fig S2C, Fig. 3B). We quantified the formation of thiol-groups upon covalent linkage of inorganic sulfide to organic backbones using Ellman’s reagent and spectrophotometry (Ellman, 1959). For the reaction of OAHS to homocysteine, detection of thiol formation was hampered by the presence of a high background, although this could be reduced upon the removal of either pyridoxal-5’phosphate (PLP) or EDTA from the reaction, suggesting a non-enzymatic cause. Despite these constraints, we observed a slight increase in thiol levels in reactions containing immuno-purified YLL058Wp (Fig. 3E). Conversely, the corresponding reaction of OAS to cysteine was unaffected by background, although enzymatic thiol formation could only be sustained when the enzyme concentration was high (RCT_100), (Fig. S3D). To test if these apparently low levels of organic sulfur metabolites were sufficient to promote yeast growth, we conducted a growth experiment including both the *met17*Δ*YLL058W*Δ and *met17*Δ*cys3*Δ mutants. Supernatants obtained from the *in vitro* OAHS reactions and OAS reactions (RCT_0, RCT_25 and RCT_100) were supplemented into the culture media of *met17*Δ, *met17*Δ*YLL058W*Δ and *met17*Δ*sir2*Δ deletion strains or the *met17*Δ*cys3* deletion strain respectively. At 72h post-inoculation, the endpoint OD_600_ of the OAHS supplemented cultures were compared against cultures directly supplemented with 0.15 mM methionine and homocysteine (Fig. 3F), revealing growth and the rescue of organosulfur auxotrophy in the supernatant-supplemented strains. Conversely, the supernatants taken from both OAHS and OAS reactions did not support growth of any tested strain in the absence of the immunoprecipitated enzyme (RCT_0) (Fig. 3C and Fig. S3E). Most significantly, although the *met17*Δ*YLL058W*Δ strain could not be rescued by NaHS supplementation (Fig.3C), growth was restored by homocysteine either via direct supplementation or via the OAHS post-reaction supernatant (Fig. 3F), whilst supplementation of the OAS post-reaction supernatant rescued the growth of the *met17*Δ*cys3*Δ strain, indicating the formation of cysteine (Fig. S3E). Hence, the reaction products formed in the presence of YLL058Wp were sufficient to rescue the growth defects of the organosulfur auxotrophs. Furthermore, we also observe that the *met17*Δ*YLL058W*Δ strain is unable to be reduced by NaHS supplementation, proving that YLL058Wp is indeed the key enzyme for sulfide utilization from H_2_S (Fig. S3F). Interestingly, in reaffirming that NaHS supplementation does not rescue growth in the absence of YLL058Wp, we also found that *met17*Δ, *met17*Δ*YLL058W*Δ and *met17*Δ*str2*Δ strains are inherently sensitive to cysteine-induced toxicity; cysteine supplementation either reduced or abolished growth in all strains (Fig. S3F). Since YLL058Wp is capable of H_2_S-dependent homocysteine synthase activity the sufficiently restores growth in the absence of Met17p, we henceforth refer to YLL058Wp as **H**ydrogen **S**ulfide **Utilizing**-1 (Hsu1p). We further observed that the OAHS post-reaction supernatant does not rescue growth to the same degree in *met17*Δ*sir2*Δ compared to the *met17*Δ*YLL058W*Δ strain, perhaps influenced by the impaired ability to detoxify cysteine to cystathionine (Fig. 3B, Fig. 3F, Fig. S3F). The *met17*Δ*cys3*Δ mutant was able to grow when NaHS is supplemented, indicating that there is an alternative route from H_2_S to cysteine that does not use cystathionine as an intermediate (Fig 3B, Fig S2C). Although it is tempting to suggest that YLL058Wp could provide a direct route from H_2_S to cysteine (Fig 3B, gray lines) as it is able to catalyze the conversion of OAS to cysteine *in vitro* this is unlikely to occur in yeast that descend from the S288c background due to a lack of a functional serine O-acetyltransferase (SAT, (Cherest & Surdin-Kerjan, 1992; B. I. Ono et al., 1999; Takagi et al., 2003)), therefore suggesting a different uncharacterised route from H_2_S to cysteine that is independent of Cys3p.

## Discussion

Sulfur assimilation is a fundamental process of life, yet the biological response to sulfur limitation is still not fully understood. This study was stimulated by an aim to solve the biochemical basis for a paradoxical yeast growth phenotype reported by Cost and Boeke (Cost & Boeke, 1996; G. J. Cost, 1999) when they established *MET17* as an auxotrophic marker for yeast genetic experiments. Certain specific conditions render this auxotrophic marker ‘leaky’; cells are somehow able to overcome an otherwise strict organosulfur auxotrophy when inoculated at high cell density. They also found that a peculiarity of *met17*Δ cells is the overproduction and leakage of sulfide ions, which we have termed sulfide overflow. In searching for the biochemical basis that unites these observations, we find evidence for a previously overlooked sulfur fixation route. This protein, herein named Hsu1p, allows cells to re-incorporate the sulfide ions that accumulate to growth-supporting concentrations in high-density cultures. Hsu1p resolves organosulfur auxotrophy by substituting for Met17p, catalyzing the incorporation of inorganic sulfide ions with OAHS to form homocysteine, thereby replenishing the central organosulfur pool and rescuing the *met17*Δ growth defect in a cell-cell cooperative manner. Our data also suggests an explanation for the temperature dependency of the rescue; higher temperatures drive the phase equilibrium towards formation of gaseous H_2_S, which is readily lost from the system.

Transcriptomic data recorded by Oliver and colleagues (Boer et al., 2003; Zhang et al., 2001) show that Hsu1p is induced during sulfur limitation, although how the expression of this additional homocysteine synthase can contribute to a growth advantage remains unclear. We observed that the accumulation of H_2_S overcomes the loss of Met17p in the presence of Hsu1p, either directly provided or generated as a result of sulfide overflow in high density cultures. Our data further indicates that the expression of Hsu1p appears to respond to high sulfide levels, which suggests Hsu1p might be a less efficient enzyme compared with Met17p and require higher concentrations of H_2_S to support growth. Therefore, it seems likely that homocysteine synthase activity may be a promiscuous reaction of Hsu1p and much like its ortholog Str2p, may primarily act as cystathionine-γ-synthase. However, the structural basis that prevents Str2p adopting the same promiscuous activity in the absence of Met17p requires further investigation. The ability of *met!7\* mutants to overcome methionine auxotrophy with Hsu1p illustrates an interesting property of feedback control in metabolism. Negative feedback is a common feature of metabolic pathways that drives upstream enzymes to be upregulated in an attempt to increase flux through the pathway when low concentrations of downstream metabolites are detected (Monod et al., 1978; Sauro, 2017; Yates & Pardee, 1956). Feedback or end-point inhibition has been shown to exist in various biosynthetic pathways in yeast including the synthesis of methionine, arginine, branched-chain amino acids, lysine and aromatic amino acids (Godard et al., 2007; Ljungdahl & Daignan-Fornier, 2012; Thomas & Surdin-Kerjan, 1997). We see that in *met17*Δ mutants, the upregulation of these upstream enzymes can lead to build up of sufficient H_2_S to allow Hsu1p to bypass the metabolic deficiency with a reaction that does not occur in its absence. Therefore, gaseous H_2_S could theoretically even function as a chemical messenger, a shared metabolite that could allow cells within the culture to act as a quorum sensing mechanism.

An interesting aspect of the Met17 conundrum is that it shows that sulfur fixation, as an essential metabolic process, can involve cell-cell metabolic cooperation. Most microbes live in microbial communities that contain auxotrophic species. In fact, from all 12,538 microbial communities sequenced as part of the Earth Microbiome project (Thompson et al., 2017), only 6 contained no amino acid auxotrophs that obtain intermediates from other communal cells (Yu et al., 2022). Metabolic interactions between individual cells are still understudied, but might prove key in understanding the collective metabolic properties of microbial communities.

## Acknowledgements

We thank all our lab members and Prof. Judith Berman for critical discussion and comments to the manuscript. We also thank Dr. Svend Kjaer and Dr. Phillip Walker at the Crick Structural Biology STP for their technical support. For the purpose of Open Access, the author has applied a CC BY public copyright license to any Author Accepted Manuscript version arising from this submission.

This work was supported by the following funding bodies:

Wellcome Trust IA grant IA 200829/Z/16/Z (MR, JSLY, CC-M, BMH, SKA, VD, CBM)

Ministry of Education and Research MSCoresys 031L0220 (MR, CTL, OL and FA)

European Commission CoBiotech project Sycolim ID#33 (MR, CTL, OL and FA)

Swiss National Science Foundation Postdoc Mobility fellowship 191052 (JH)

European Research Council grant ERC-SyG-202 951475 (MR)

The Francis Crick Institute, which receives its core funding from:

Cancer Research UK (FC001134)

UK Medical Research Council (FC001134)

Wellcome Trust (FC001134)

## Author contributions

Conceptualization: JSLY, MR

Methodology: VD, CBM, MM

Investigation: CCM, JH, BMH, OL, SA, AL, FA, CTL

Formal analysis: JSLY, CCM, BMH, OL, SA.

Visualization: JSLY, CCM, MR

Funding acquisition: MR

Project administration: JSLY, MR

Supervision: MR

Writing - original draft: JSLY, BMH, MR

Writing - review & editing: JSLY, BMH, MR

## Competing interests

Authors declare that they have no competing interests.

## Data materials and availability

All data are available within the main text or the supplementary materials. The proteomic dataset used for the re-analysis of the *met17*Δ phenotype can be accessed at the Proteomics Identification Database (PRIDE) with the dataset identifier PXD031160.

## Supplementary Materials

### Materials and Methods

#### Yeast culture

Yeast strains (Table S1) were streaked from cryostock onto YPD or SM+HLUM and incubated for 3 days at 30°C before a single colony was picked to inoculate 20 ml of media in a conical flask and subsequently incubated at 30°C for growth under various conditions as outlined in the experiments above. Prototrophic and SeMeCo strains were revived onto synthetic minimal (SM) media agar (6.8 g/L Yeast Nitrogen Broth without amino acids (YNB, Sigma, Y0626; Lot: MKCF2863), 2% w/v glucose (Sigma), 2% w/v agarose (Life Technologies), whilst strains harboring only *met17*Δ were streaked onto SM+Met. Strains where auxotrophy complementation or xenogeneic expression was introduced were cultured in selective media at all times unless otherwise stated. Cultures in which required activation of the bypass reaction (high-density inoculations or supplementation with NaHS) were cultured at 25°C to maximize growth (see Fig 1C.)

#### Systematic growth analysis of *met17*Δ in minimal media

BY4741 HIS3,LEU2,URA3,*met17*Δ and the prototrophic control strain were streaked on SM+Met (0.02g/l), and grown for 36 hours. Single colonies (n=3 replicates per genotype) were inoculated in SM+Met, grown o/n at 30°C in Erlenmeyer flasks, diluted 1:20 in fresh SM+Met media, and incubated for 4 hours. Cells were collected, washed in autoclaved H_2_O (Millipore), re-suspended in SM, and the OD_600_ of each strain was adjusted to 0.6. Each set of replicates (n=3 for the deletion strain, and n=3 for the control strain) were then transferred in a deep-96-well plate where a sequential 1:1 dilution of 100 μl culture was performed with 100 μl SM media to obtain an OD_600_ range of 0.3 to ~0.0003. 4×200 μl of each well was transferred to 4×96-well microtiter plates (Costar). The plates were sealed using a non-permeable, clear foil (Roth) and were incubated at 22°C, 26°C, 30°C and 37°C without shaking. Optical density was recorded by placing the plates in an Infinite M200 Pro plate reader (Tecan), and OD600 was obtained from the median of 9 multi-well reads per well. Subsequently, for each plate, the median OD600 of all blank (n=8 per plate) values was subtracted from obtained OD_600_ values.

#### Hydrogen sulfide production quantification

Qualitative assessment of hydrogen sulfide production was indicated by the precipitation of lead sulfate in growth media containing 0.1 g/L lead(II) acetate. Quantification of hydrogen sulfide was conducted using 7-azido-4-methylcoumarin (AzMC), which undergoes selective reduction of the azide moiety in the presence of hydrogen sulfide to form 7-amino-4-methylcoumarin (AMC). In brief, 10 ul of filtered yeast media was mixed with 200 ul of 10 μM AzMC and incubated at RT for 30 min before the fluorescence was recorded on a Tecan spectrophotometer at _em_λ = 450 nm (_ex_λ = 365 nm). For the generation of hydrogen sulfide, we utilized sodium hydrosulfide (NaHS) as the donor, reconstituted with water to a stock concentration of 20 mM and a working concentration in the media of 0.2-0.4 mM.

#### Isotopic tracing by LC-MS/MS

##### Sample preparation

Samples were essentially treated as described above. Briefly, single colonies were re-suspended in sterilized H_2_O and OD_600_ was determined. For each biological replicate, 2x 1.1ml of 1.7 g/l YNB without ammonium sulfate and amino acids (Sigma), 2% glucose, and naturally labeled ammonium sulfate (5 g/l) or ammonium sulfate-^34^S (5 g/L, Sigma, >= 98% ^34^S) as situated in 2x 1.5 ml microcentrifuge tubes were inoculated to an OD_600_ of 0.03. Cells were incubated in closed vials at 30 °C without shaking for 48 hours, and harvested by centrifugation (3 min, 3220 xg). The supernatant was taken off and passed through a syringe filter (0.22 μm). Corresponding pellets were pooled and extracted in 100 μl pre-warmed ethanol, vortexing, incubation for 2 min in a water bath at 80 °C, vortexed, and transferred again for 2 min to a water bath at 80 °C. Extract was cleared by centrifugation (15 min, 16000 xg at 4 °C) and the supernatant was transferred to HPLC glass vials for analysis. The filtered supernatant of each tube (0.9 ml) was reconstituted by addition of fresh media constituents as supplied by addition of 50 μl glucose (40% v/v), and 100 μl YNB w/o ammonium sulfate and amino acids (17 g/l) and respective ammonium sulfate (50 g/l). Media was then inoculated to an OD_600_ of 0.003, grown for 48 hours, and harvested and extracted as described above.

##### Data acquisition

Data was acquired by LC-MS/MS using a high pressure liquid chromatography system (1290 Infinity, Agilent Technologies) coupled to an Triple Quad mass spectrometer (6470, Agilent Technologies). Samples were separated using a Acquity UPLC BEH Amide 1.7 um, 2.1×100 mm column. The gradient consisted of mobile phase A (water with 100 mM ammonium carbonate) ramped against mobile phase B (acetonitrile) with a flow rate of 0.3ml as follows: 0 min, 70% B, 3 min, 70% B, 7 min, 40% B, 8 min, 40% B, 8.1 min, 70% B, followed by 1.9 min equilibration. Mass spectrometer was operating in MRM mode with the following source settings: gas temperature 200 C, Gas flow 13 L/min, nebulizer 60 psi, sheath gas temp 300 °C, sheath gas flow 11 L/min, Capillary Voltage 3500 (positive mode) or 2500 (negative mode) and a delta EMV of 100 (both modes). Injection volume was 1 μl.

Metabolites were identified by matching retention time and transitions as confirmed by analytical standards. Methionine was analyzed in positive mode using the transitions 150.1 -> 132.9 (unlabeled) or 152.1 -> 134.9 (^34^S labeled). Reduced glutathione was analyzed in negative mode with the transitions 306 -> 143 (unlabeled) and 308 -> 143 (^34^S labeled).

#### Immunoprecipitation of YLL058W-GFP

The strain identity and coding regions were first confirmed via specific primers commercially available, designed in-house or obtained from the original study ((Winzeler et al., 1999), CHK: 5’-CTTCAGGAGCGCTAATTTCG-3’, F2CHK: 5’-AACCCGGGGATCCGTCGACC-3’, GFP-F: 5’-GGTCCTTCTTGAGTTTGTAAC-3’, GFP-R: 5’ -CCATCTAATTCAACAAGAATTGGGACAAC-3’, pHybLex-R: 5’-GAGTCACTTTAAAATTTGTATACAC-3). To maximize the recovery of YLL058W-GFP protein, *BY4741*(*MATa, his3*Δ*1*, *leu2*Δ, *ura3*Δ, *met17*Δ*YLL058w::YLL058W-GFP-HIS3MX6*) grown from the GFP library was cultured for 5 days in 150 ml minimal media supplemented with leucine, uracil and 0.4 mM NaHS at 25 °C. Following this, cells were pelleted by centrifugation at 3220 x*g* for 5 min at 4 °C and resuspended in an appropriate volume of Y-PER™ Y east Protein Extraction Reagent (ThermoFisher) containing 1x Protease Inhibitor cocktail (P8215, Sigma). Purified protein extract was obtained following the manufacturer’s protocol, to which 250 ul of GFP-clamp resin (in-house, Crick-Structural Biology Scientific Technology Platform) was added to the lysis solution and incubated at 4 °C for 2 h with rotation.

#### Hydrogen sulfide utilization enzyme and growth assays

Following immunoprecipitation, beads were washed with twice with Y-PER™ Y east Protein Extraction Reagent (ThermoFisher), followed by a final wash with reaction buffer (0.1 M potassium phosphate buffer pH 7.8, 1 mM ethylenediaminetetraacetic acid (EDTA), 200 μM pyridoxal 5’-phosphate (PLP), all Sigma). Enzyme-bead complexes were then re-suspended in 100 μl reaction buffer containing 5 mM O-acetyl-serine (OAS, Sigma) or O-acetyl-homoserine (OAHS, Insight Biotech) and placed into Eppendorf tube holders contained in a large beaker. A smaller beaker containing 20 mM NaHS solution was also placed into the large beaker which was sealed in parafilm and made airtight. The entire setup was placed at 30 °C and incubated for 24 h, during which H_2_S would out-gas from the NaHS solution, dissolve and react with OAS/OAHS in the presence of the enzyme (Fig. S3C). Tubes were then retrieved from the beaker and placed in an Eppendorf Thermomixer and shaken for 2 h at RT to remove residual H_2_S before quantification via Ellman’s reagent (ThermoFisher). For growth assays, 50 μl of reaction supernatant was transferred to 15 ml round bottom tubes containing 3 ml of media and incubated with shaking for up to 5 days at 25°C before endpoint OD_600_ was measured using a cuvette spectrophotometer.

#### Quantification of cysteine/homocysteine

Quantification of cysteine/homocysteine resulting from the OAS/OAHS reaction respectively was performed using Ellman’s reagent (5,5-dithio-bis-(2-nitrobenzoic acid, ThermoFisher, (Ellman, 1959)), which preferentially reacts with sulfhydryl groups to yield a colored product. In brief, a standard curve using homocysteine (0-0.2 mM, Sigma) was generated and incubated with the reagent following the manufacturer’s protocol at a 1:1 ratio. Absorbance was measured on a Tecan Infinite M200 PRO® spectrophotometer at λ = 412 nm and concentration was calculated using the standard curve.

## Supplementary Figures

**Supplementary figure 1:**
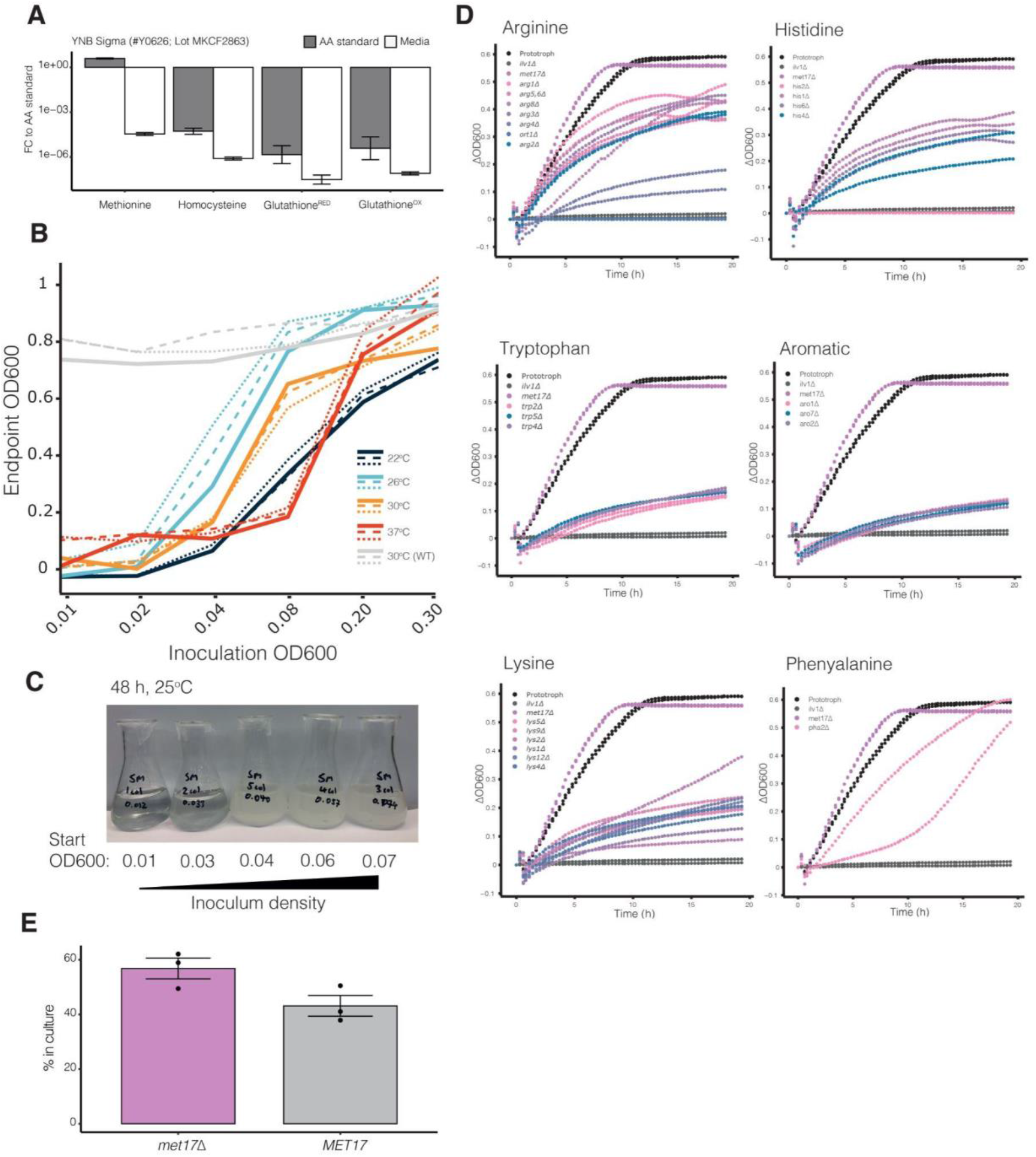
Characterisation of the *met17*Δ prototrophic growth phenotype in the absence of methionine. **(A)** LC-MS analysis of Yeast Nitrogen Broth (YNB) component of our minimal media. Quantified metabolites were compared to their respective standards and fold-change (FC) calculated. GSH: glutathione, GSSG: reduced glutathione. **(B)** Liquid cultures of *met17*Δ strain in minimal media without methionine inoculated at 6 different cell densities (0.01 to 0.3) and cultured at 4 different temperatures (22, 26, 30, 37°C) for 24h. Gray indicates wild-type control strain that is prototrophic for methionine cultured at 30°C. Lines indicate three replicates per condition, where n=3. **(C)** Growth of *met17*Δ strain in 20 ml cultures. Starting OD_600_ of cultures varied between 0.01 to 0.07 and growth was assessed after 48h of culture in minimal media without methionine. **(D)** Growth curves captured from indicated amino acid auxotrophic strains compared against the prototrophic and *ilvΔ* strains. **(E)** Quantification of methionine auxotrophs in exponentially growing SeMeCo cultures. Data are mean±SEM from 3 biologically independent replicates (n=3),

**Supplementary figure 2:**
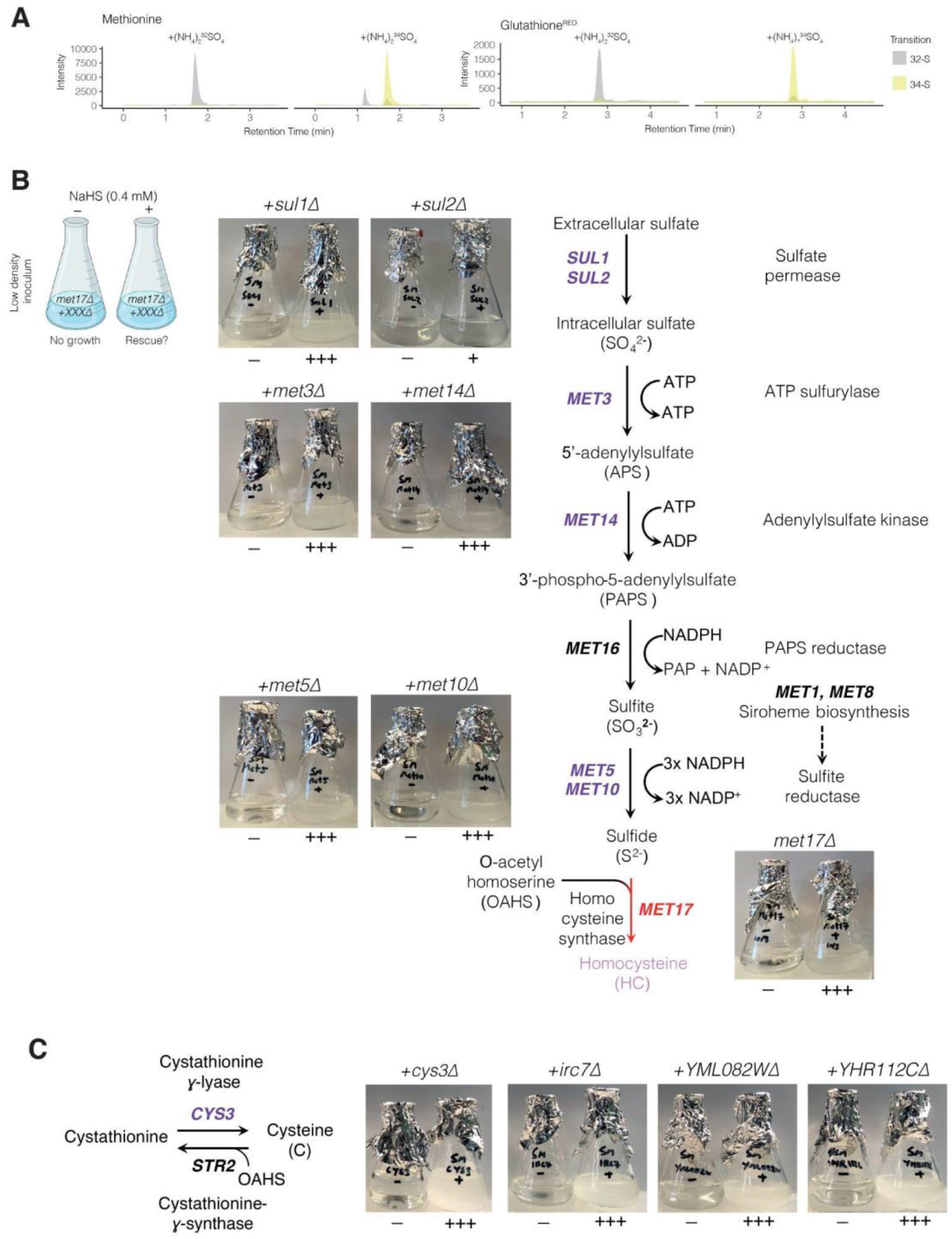
Cell density dependency and involvement of sulfur assimilation mutants in the utilization of H_2_S. **(A)** Representative mass spectroscopy traces of methionine, reduced glutathione in cultures supplemented with ^32^S and ^34^S-labeled ammonium sulfate. Gray and yellow traces indicate S^32^ and S^34^ versions of the metabolite. **(B)** Growth screens for loss of H_2_S utilization in sulfur assimilation deletion mutants upstream of *MET17* mapped against the pathway. **(C)** Screen for loss of H_2_S utilization in strains lacking the Met17p orthologs Cys3p, Irc7p, YML082Wp, and YHR112Cp. All mutants tested are in a background of *met17Δ.* Scheme for Fig. SB was created with BioRender.com.

**Supplementary figure 3:**
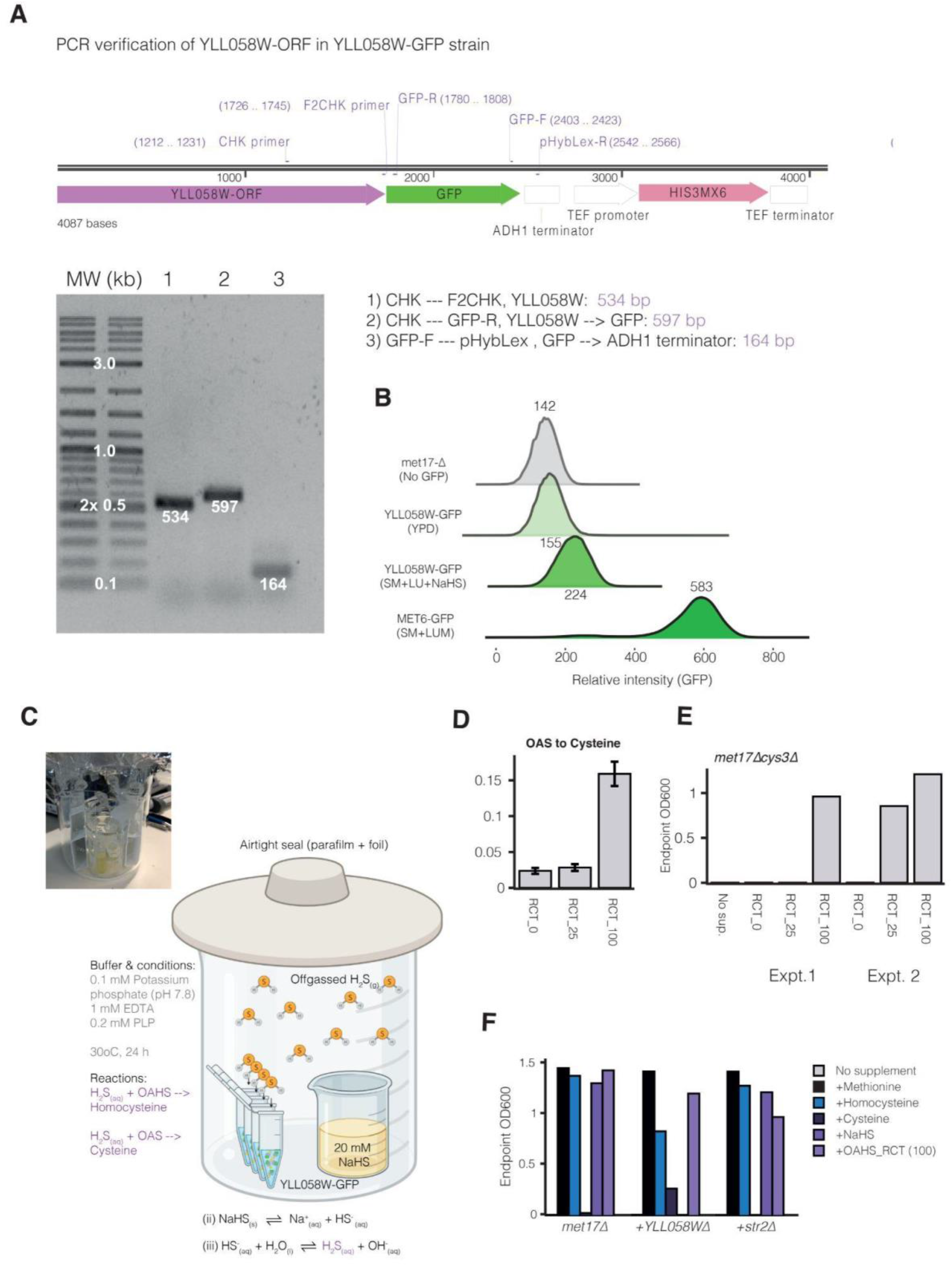
Characterization of YLL058W expression and enzymatic activities. **(A)** PCR verification of the YLL058W-GFP ORF using genomic DNA isolated from the YLL058W-GFP strain using primers within the YLL058W and GFP open reading frames and the ADH1 terminator. All products were of the expected size as indicated by gel electrophoresis. **(B)** Flow cytometry analysis and confirmation of GFP expression intensity in response to histidine and methionine deficiency with H_2_S supplementation. Numbers indicate median fluorescence intensity, from ~40,000 events captured per strain. The *met17*Δ strain was used as a negative, non-fluorescent control, whilst the Met6p-GFP strain was used as a positive, highly fluorescent control. (**C**) Schematic of H_2_S utilization enzyme assay. Off-gassing of H_2_S from a 20 mM NaHS solution dissolves into reaction tubes containing either OAHS or OAS as substrate. Insert indicates actual setup of reactions **(D)** Quantification of thiol concentrations via Ellman’s reagent using supernatant from enzyme assays where immunoprecipitated YLL058W-GFP was incubated with either OAS as the organic substrate. RCT_0, 25 and 100 indicate volumes of enzyme-resin slurry used in the reaction. Yield = [product]/[substrate], OAS: 0.16/5 = 3.2%. Data are mean thiol concentration ± SD where n = 3 biologically independent replicates. **(E)** Growth rescue assay of *met17*Δ*cys3*Δ strain by product of the OAS reaction. RCT_0, 25 and 100 indicate the volume of enzyme-resin complex used which approximates increasing enzyme concentrations. OD_600_ was measured from two independent sets of experiments as indicated. (**F**) Endpoint OD_600_ measurements of *met17*Δ and double deletion strains (*met17*Δ, +*YLL058W*Δ, and +*str2*Δ) supplemented with various sulfur sources (methionine, cysteine, homocysteine at 0.15 mM, NaHS at 0.4 mM). OAHS_RCT indicates supplementation with supernatant from *in vitro* enzyme assay with OAHS as substrate. Data is from one independent experiment, but replicate observations in Fig 2F. Scheme for Fig. 3C was created with BioRender.com.

**Supplementary Table 1:** Yeast strains used in this study.

| Name | Source | Genotype | Experiment used |
| --- | --- | --- | --- |
| Prototrophic | Yu et al., 2022 | *HIS3,LEU2,URA3,MET17* | Fig. 1B-C, 1E, S1D |
| *met17Δ* | Yu et al., 2022 | *HIS3,LEU2,URA3,met17Δ* | Fig. 1B-D, 1F-H, S1B-C, S3F, 2A-F, |
| 4p-SeMeCo | Yu et al., 2022 | *pHIS3,pLEU2,pURA3,pMET17* | Fig. 1E, S1E |
| pM-CFP-SeMeCo | Yu et al., 2022 | *HIS3,LEU2,URA3,pMET17:CFP* | Fig. 1F-H |
| *met17Δ*<br>(aux-library) | Winzler et al., 1999 | *his3Δ,leu2Δ,ura3Δ,met17Δ* | Fig. 1D, S1D, 3A, 3F |
| *ilv1Δ* | Winzler et al., 1999 | *his3Δ,leu2Δ,ura3Δ,met17Δ,ilv1Δ* | Fig. 1D, S1D |
| *met5Δ* | Winzler et al., 1999 | *his3Δ,leu2Δ,ura3Δ,met17Δ,met5Δ* | Fig. 1D |
| *met3Δ* | Winzler et al., 1999 | *his3Δ,leu2Δ,ura3Δ,met17Δ,met3Δ* | Fig. 1D |
| *met1Δ* | Winzler et al., 1999 | *his3Δ,leu2Δ,ura3Δ,met17Δ,met1Δ* | Fig. 1D |
| *met7Δ* | Winzler et al., 1999 | *his3Δ,leu2Δ,ura3Δ,met17Δ,met7Δ* | Fig. 1D |
| *arg1Δ* | Winzler et al., 1999 | *his3Δ,leu2Δ,ura3Δ,met17Δ,arg1Δ* | Fig. S1D |
| *arg5,6Δ* | Winzler et al., 1999 | *his3Δ,leu2Δ,ura3Δ,met17Δ,arg5,6Δ* | Fig. S1D |
| *arg8Δ* | Winzler et al., 1999 | *his3Δ,leu2Δ,ura3Δ,met17Δ,arg8Δ* | Fig. S1D |
| *arg3Δ* | Winzler et al., 1999 | *his3Δ,leu2Δ,ura3Δ,met17Δ,arg3Δ* | Fig. S1D |
| *arg4Δ* | Winzler et al., 1999 | *his3Δ,leu2Δ,ura3Δ,met17Δ,arg4Δ* | Fig. S1D |
| *arg2Δ* | Winzler et al., 1999 | *his3Δ,leu2Δ,ura3Δ,met17Δ,arg2Δ* | Fig. S1D |
| *ort1Δ* | Winzler et al., 1999 | *his3Δ,leu2Δ,ura3Δ,met17Δ,ort1Δ* | Fig. S1D |
| *his2Δ* | Winzler et al., 1999 | *his3Δ,leu2Δ,ura3Δ,met17Δ,his2Δ* | Fig. S1D |
| *his1Δ* | Winzler et al., 1999 | *his3Δ,leu2Δ,ura3Δ,met17Δ,his1Δ* | Fig. S1D |
| *his6Δ* | Winzler et al., 1999 | *his3Δ,leu2Δ,ura3Δ,met17Δ,his6Δ* | Fig. S1D |
| *his4Δ* | Winzler et al., 1999 | *his3Δ,leu2Δ,ura3Δ,met17Δ,his4Δ* | Fig. S1D |
| *trp2Δ* | Winzler et al., 1999 | *his3Δ,leu2Δ,ura3Δ,met17Δ,trp2Δ* | Fig. S1D |
| *trp4Δ* | Winzler et al., 1999 | *his3Δ,leu2Δ,ura3Δ,met17Δ,trp4Δ* | Fig. S1D |
| *trp5Δ* | Winzler et al., 1999 | *his3Δ,leu2Δ,ura3Δ,met17Δ,trp5Δ* | Fig. S1D |
| *aro1Δ* | Winzler et al., 1999 | *his3Δ,leu2Δ,ura3Δ,met17Δ,aro1Δ* | Fig. S1D |
| *aro2Δ* | Winzler et al., 1999 | *his3Δ,leu2Δ,ura3Δ,met17Δ,aro2Δ* | Fig. S1D |
| *aro7Δ* | Winzler et al., 1999 | *his3Δ,leu2Δ,ura3Δ,met17Δ,aro7Δ* | Fig. S1D |
| *lys5Δ* | Winzler et al., 1999 | *his3Δ,leu2Δ,ura3Δ,met17Δ,lys5Δ* | Fig. S1D |
| *lys9Δ* | Winzler et al., 1999 | *his3Δ,leu2Δ,ura3Δ,met17Δ,lys9Δ* | Fig. S1D |
| *lys2Δ* | Winzler et al., 1999 | *his3Δ,leu2Δ,ura3Δ,met17Δ,lys2Δ* | Fig. S1D |
| *lys1Δ* | Winzler et al., 1999 | *his3Δ,leu2Δ,ura3Δ,met17Δ,lys1Δ* | Fig. S1D |
| *lys12Δ* | Winzler et al., 1999 | *his3Δ,leu2Δ,ura3Δ,met17Δ,lys12Δ* | Fig. S1D |
| *lys4Δ* | Winzler et al., 1999 | *his3Δ,leu2Δ,ura3Δ,met17Δ,lys4Δ* | Fig. S1D |
| *pha2Δ* | Winzler et al., 1999 | *his3Δ,leu2Δ,ura3Δ,met17Δ,pha2Δ* | Fig. S1D |
| *cys3Δ* | Winzler et al., 1999 | *his3Δ,leu2Δ,ura3Δ,met17Δ,cys3Δ* | Fig. S2C, S3E |
| *met6Δ* | Winzler et al., 1999 | *his3Δ,leu2Δ,ura3Δ,met17Δ,met6Δ* | Fig. 3A |
| *met2Δ* | Winzler et al., 1999 | *his3Δ,leu2Δ,ura3Δ,met17Δ,met2Δ* | Fig. 3A |
| *str2Δ* | Winzler et al., 1999 | *his3Δ,leu2Δ,ura3Δ,met17Δ,str2Δ* | Fig. 3A, 3F,<br>S3F |
| *str3Δ* | Winzler et al., 1999 | *his3Δ,leu2Δ,ura3Δ,met17Δ,str3Δ* | Fig. 3A |
| *YLL058WΔ/Hsu1Δ* | Winzler et al., 1999 | *his3Δ,leu2Δ,ura3Δ,met17Δ,hsu1Δ* | Fig. 3A, 3F,<br>S3F |
| *sul1Δ* | Winzler et al., 1999 | *his3Δ,leu2Δ,ura3Δ,met17Δ,sul1Δ* | Fig. S2C |
| *sul2Δ* | Winzler et al., 1999 | *his3Δ,leu2Δ,ura3Δ,met17Δ,sul2Δ* | Fig. S2C |
| *met14Δ* | Winzler et al., 1999 | *his3Δ,leu2Δ,ura3Δ,met17Δ,met14Δ* | Fig. S2C |
| *met10Δ* | Winzler et al., 1999 | *his3Δ,leu2Δ,ura3Δ,met17Δ,met10Δ* | Fig. S2C |
| *irc7*Δ | Winzler et al., 1999 | *his3*Δ,*leu2*Δ,*ura3*Δ,*met1*7Δ,*irc7*Δ | Fig. S2C |
| *YML082C*Δ | Winzler et al., 1999 | *his3*Δ,*leu2*Δ,*ura3*Δ,*met1*7Δ,*YML082C*Δ | Fig. S2C |
| *YHR112C*Δ | Winzler et al., 1999 | *his3*Δ,*leu2*Δ,*ura3*Δ,*met1*7Δ,*YHR112C*Δ | Fig. S2C |
| *YLL058W*-GFP/<br>*Hsu1*-GFP | Huh et al., 2003 | *leu2*Δ,*ura3*Δ,*met1*7Δ,*HSU1*-<br>*GFP::HisMX6* | Fig. S3B, 4B<br>S4 |
| *MET6*-GFP | Huh et al., 2003 | *leu2*Δ,*ura3*Δ,*met1*7Δ,*MET6*-<br>*GFP::HisMX6* | Fig. S3B |

## Notes

https://www.ebi.ac.uk/pride/archive/projects/PXD031160

### Competing Interest Statement

The authors have declared no competing interest.

